# Evidence of phylosymbiosis in *Formica* ants

**DOI:** 10.1101/2023.01.20.524858

**Authors:** Raphaella Jackson, Patapios A. Patapiou, Gemma Golding, Heikki Helanterä, Chloe K. Economou, Michel Chapuisat, Lee M. Henry

**Author notes:** **Correspondence:** Lee M. Henry.

## Abstract

Insects share intimate relationships with microbes that play important roles in their biology. Yet our understanding of how host-bound microbial communities assemble and perpetuate over evolutionary time is limited. Ants host a wide range of microbes with diverse functions and are an emerging model for studying the evolution of insect microbiomes. Here, we ask whether phylogenetically related ant species have formed distinct and stable microbiomes. To answer this question, we investigated the microbial communities associated with queens of 14 *Formica* species from five clades, using deep coverage 16S rRNA amplicon sequencing. We reveal that *Formica* species and clades harbour highly defined microbial communities that are dominated by four bacteria genera: *Wolbachia, Lactobacillus*, *Liliensternia,* and *Spiroplasma*. Our analysis reveals that the composition of *Formica* microbiomes mirrors the phylogeny of the host, i.e. phylosymbiosis, in that related hosts harbour more similar microbial communities. Our analysis also revealed significant correlations between microbe co-occurrences, which suggests that synergistic and antagonistic interactions may contribute to the phylosymbiotic signal. Additional factors potentially contributing to the phylosymbiotic signal are discussed, including host phylogenetic relatedness, host-microbe genetic compatibility, modes of transmission, and similarities in host ecologies (e.g., diets). Overall, our results support the growing body of evidence that microbial community composition closely depends on the phylogeny of their hosts, despite bacteria having diverse modes of transmission and localisation within the host.

## Introduction

All multicellular organisms interact with microbes that influence their biology, and this is particularly true for insects. Microbial associates of insects are integral to many aspects of their hosts’ biology, including nutrition, immunity, reproduction, growth, and metabolism (Feldhaar et al., 2007; Horak et al., 2020; Singh and Linksvayer, 2020). Both beneficial and parasitic relationships between insects and their microbes can therefore drive important population processes such as speciation and ecological expansion, by unlocking previously inaccessible niches (Russell et al., 2009; Bennett and Moran, 2015; Sanders et al., 2017). Yet, we are only beginning to understand how the communities of microbes associated with insects – their microbiomes - assemble and perpetuate over evolutionary time.

Ants are an excellent system for understanding how and why microbiomes evolve, as they have formed a wide range of relationships with microbes, from stable associations with a single microbe to relatively complex gut communities. Stable associations include heritable endosymbionts, such as *Wolbachia*, *Arsenophonus*, and *Spiroplasma* (Russell et al., 2012), as well as *Blochmannia* and *Westeberhardia*, the ancient bacteriocyte-associated symbionts found in *Camponotus* and *Cardiocondyla* ants, respectively (Degnan et al., 2002; Klein et al., 2016). Several ant genera have also evolved stable communities of microbes housed in their guts, such as *Cephalotes* turtle ants, whereas other ant lineages show little to no consistent associations with any gut-associated microbes (Sanders et al., 2017; Hu et al., 2018). While there are a few detailed investigations of unique, vertically transmitted symbionts (e.g. Jackson et al., 2022, Klein et al., 2016), and broad comparative analyses across phylogenetically diverse host species (e.g. Sanders et al., 2017, Russell et al., 2009), ants, in general, have been understudied for microbes. In particular, few studies have investigated what shapes the community of microbes associated with clades of phylogenetically related ant species.

Studying microbiome structure across related hosts helps us disentangle the factors driving patterns observed in individual species and answer key questions, such as: whether phylogenetically related hosts evolved similar microbiomes, a signature of phylosymbiosis (Lim and Bordenstein, 2020); whether the presence of host-bound microbes is influenced by antagonistic or synergistic interactions with other microbes; or whether shared host ecologies shape the communities of microbes insects carry (Brooks et al., 2017). *Formica* ants are an ideal model to address these questions, as they have formed persistent associations with at least two heritable symbionts, the bacteriocyte-associated mutualist *Liliensternia*, and the reproductive parasite *Wolbachia*, as well as with gut-associated microbes such as *Lactobacillus* (Russell, 2012; Jackson et al., 2022; Zheng et al., 2022).

Here, we ask whether *Formica* ants harbour stable communities of microbes. To answer this question, we assess the composition and consistency of microbiomes in queens from 14 *Formica* species belonging to five clades, using deep coverage 16S rRNA amplicon sequencing. We focus the analysis on queens, because ant workers have been found to lose ovarially transmitted microbes as they age (Wenseleers et al., 2002; Wolschin et al., 2004). We first identify the genera of microbes associated with *Formica* ants. We then test whether there are significant differences in microbiome composition, and what microbes make up these differences, across *Formica* species and clades. Based on these results, we then test whether the microbiome composition of *Formica* species has a phylogenetic signal by examining if phylogenetically related hosts are more likely to carry similar microbial communities, and test for potential antagonistic or synergistic interaction between microbes.

## Materials and Methods

### Sample Collection Procedure

Samples were collected by opening or digging up ant colonies, sorting workers and queens by hand, and preserving them in ethanol. The location of each sample collection and identity of the collector are listed in Supplementary Table S1.

### 16S rRNA Sequencing Procedure

DNA was extracted from whole bodies of adult queens. We screened 130 adult queens from 89 colonies across 14 species, using two separate runs of 16S rRNA sequencing. See Table S1 for samples identified by run. Samples were acquired at different times and were therefore sequenced on two separate Illumina runs. We included negative controls on each run.

In run 1, we used the 515F/806R primer pair (Caporaso et al., 2011) to amplify the V4 region of the 16S rRNA gene (Table S6). All PCR reactions were performed using Q5 High-Fidelity master mix (New England Biolabs, Ipswich, Massachusetts, USA). For the first stage PCR, amplification conditions were as follows: initial denaturation at 98°C for 30 s followed by 25 cycles of 98 °C for 10 s, 50 °C for 15 s, 72 °C for 20 s and a final extension of 72 °C for 5 min. PCR clean-ups were performed using AMPureXP beads (Beckman Coulter Life Sciences, Indianapolis, United States) and then a second stage PCR was carried out to attach dual indices and Illumina sequencing adapters. Second stage PCR conditions were as follows: 95 °C for 3 min followed by 8 cycles of 98 °C for 20 s, 55 °C for 15 s, 72 °C for 15 s and a final extension of 72 °C for 5 min. A second PCR clean-up using AMPure XP beads was performed to clean up the libraries before quantification. Individual PCR products were quantified using the Qubit dsDNA HS Assay Kit (Thermo Fisher Scientific, Massachusetts, United States) and the libraries were then normalized and pooled. The pool was sequenced at Edinburgh Genomics (University of Edinburgh) on an Illumina MiSeq (paired-end, 2 x 250 bp reads).

For run 2, we used the same primer pair, PCR mix and PCR conditions as in run 1, however the PCR products were submitted to the Centre for Genomic Research (University of Liverpool) for addition of indices and adapters, and pooling of libraries. Sequencing was then carried out on a MiSeq (paired-end, 2 x 250 bp reads).

### Creation of Amplicon Sequence Variants (ASVs) and Operational Taxonomic Units (OTUs)

Adaptor sequences were removed using the function “ILLUMINACLIP” of Trimmomatic V.0.38 (Bolger et al., 2014), under default parameters. Each sequencing run was then imported into the Qiime2 bioinformatics platform (Bolyen et al., 2019) and denoised into ASVs using DADA2 (Callahan et al., 2016). We then filtered the resulting ASVs for potential contamination by discarding those at <1% relative abundance within a sample and removed ASV’s belonging to mitochondria, chloroplasts, and non-bacterial taxa. Trace ASVs unique to controls were removed as potential contaminants, as were ASVs in negative controls that were also ubiquitously found in all other samples within a sequencing run. ASVs at trace levels in controls that were also present in other samples at higher abundances, but not ubiquitous across samples, were retained. After filtering, all samples with a summed count of < 3000 ASVs were removed which accounted for all controls and two experimental samples. This filter was based on examining the total count of ASVs per sample and identifying a cut-off point based on a natural separation of the data (Supplementary Image SI1). We then ensured there was no significant relationship between total number of ASVs observed and number of unique ASVs observed (Supplementary Image SI2). For each sequencing run, filtered ASVs were then combined into a single table (Table S6). We then clustered ASVs into 97% OTUs using Vsearch (Rognes et al., 2016). ASV and OTU sequences were assigned to taxons using Qiime2’s classification algorithm with the silva 138 99% classifier database. ASV and OTU tables, alongside taxonomic assignments, were then exported for additional analysis in R Statistical Software v4.1.2 (R Core Team, 2019). All statistical analyses were conducted on ASVs. OTU data was used for visualisation purposes only, and was not included in statistical analysed.

### Statistical Analyses and Data Visualization

In R, data were normalized by changing ASV counts per sample into a percentage of total ASVs for each sample (relative abundance). To create a visualization of the ASV data, ASVs were collapsed to the genus level. To visualize the data, any genus present in more than one sample, or >10% relative abundance in a single sample, was given a distinct colour. All other genera were represented in grey as “Other”. Bar graphs depicting the relative abundance of the dominant bacterial genera within each sample were then produced using ggplot2 (Wickham, 2016).

For analysing differences in bacterial community composition, all ASV data was converted into a Bray-Curtis distance matrix using the vegdist function from the vegan package in R (Oksanen et al., 2013). A permanova analysis of the distance matrix was performed using adonis. The first comparison made was DistanceMatrix ~ Sequencing Run, to investigate the impact of Sequencing Run on the centroids and dispersion of samples. We then investigated the effect of clade and species with run as both an interactive effect (DistanceMatrix~ Run*Clade/Species) and as a blocked effect (DistanceMatrix~Clade/Species, strata=Run). Clade was defined in a manner identical to Romiguier et al. (2018). All analyses were run for 1000 permutations. We analysed the beta-dispersion using ANOVA and betadisper for both species and clade level differences. SIMPER tests were also run for species and clade, using 100 permutations. Finally, a meta-NMDS visualization was created using the metaMDS function in combination with ggplot2.

### Co-occurrence analysis

To test patterns of co-occurrence between genera of ASVs, we used a Fisher’s exact test on the count of samples with genus A only, genus B only, genus A and B, and neither genus. Fisher exact tests were performed on ASVs collapsed by bacterial genus under the following conditions: i) only bacterial genera found in the same *Formica* species and run were compared; ii) only samples from these shared run/species combination were counted in statistical comparisons, we excluded samples from run/species combinations where only one bacteria occurred iii) only bacterial genera found in >2 samples were considered in the analysis. A significance level of p<0.05 was considered significant for the Fisher’s exact test. All data for each significant analysis is available in Supplementary Table S4.

### Ribotype Diversity Visualization

To visualise the distribution of bacterial ribotypes across host species, we used 97% OTUs identified as being either *Lactobacillus*, *Wolbachia*, *Spiroplasma*, or *Liliensternia* to reconstruct a tree using the Silva ACT (Alignment, Classification and Tree) service (Pruesse et al., 2012; Quast et al., 2012) (Fig. S1). The workflow used was “add to neighbours tree” with RaxML, GTR, and GAMMA as additional parameters. Minimal identity with query sequence was set to 97% and maximum matches to query sequence were 2. The complete tree is available in Fig. S1, while Fig. 3 shows a pruned version of the tree. ITOL (Letunic and Bork, 2021) was used to visualize this pruned tree and the relative abundance data from the relevant OTUs.

### Phylobiome Creation Procedure

To generate phylobiome trees (microbiome phylogenies), we followed the procedure outlined by Brooks et al. (2017) and Lim and Bordenstein (2020). Individuals belonging to the same species were collapsed into single columns in the ASV table. ASV tables containing a single symbiont of interest (e.g., *Lactobacillus*, *Wolbachia*) were also created. We then created distance-matrices using the Qiime function “diversity beta-rarefaction” for the full collapsed ASV table and the ASV table of each symbiont of interest. We used the Bray-Curtis distance metric and the upgma clustering method. Sampling depth was 3000 and the process was repeated for 100 iterations. Phylogenies were visualized using ITOL (Letunic and Bork, 2021).

We used TreeCmp (Goluch et al., 2020) to test for congruence between the phylobiome tree and host phylogeny. TreeCmp generated scores normalized (to Unifrac) Robinson-Folds Cluster and Matching Cluster scores for each tree comparison. To determine p-values for these scores we preformed the same congruence testing for 1000 randomly generated trees. Random phylogenies were generated using T-Rex (Alix et al., 2012).

## Results

### Four Common Bacterial Genera are Associated with *Formica* Ants

We used 16S rRNA amplicon sequence variants (ASVs), taxonomically classified at genus, to quantify the composition of the microbiome of queens from 14 *Formica* species from five clades (Fig. 1, Table S1). We used this information to evaluate the presence and abundance of the known symbionts of *Formica* ants, *Liliensternia*, *Wolbachia* and *Lactobacillus*, and to identify other commonly occurring bacterial genera.

**Figure 1:**
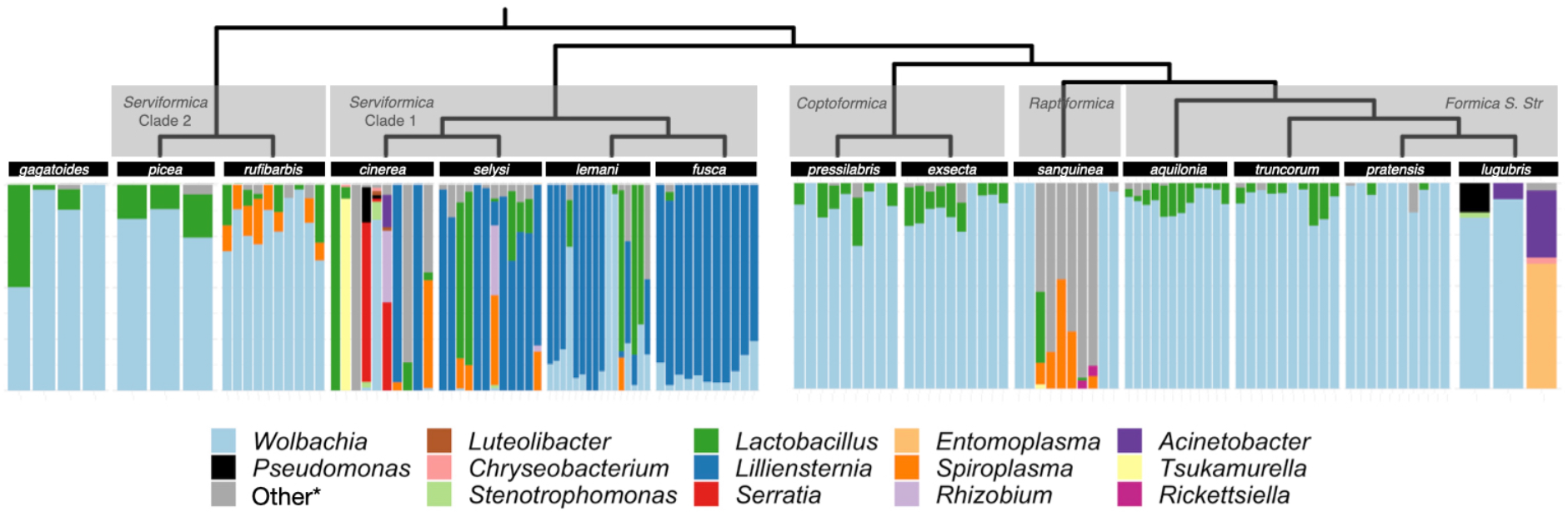
Microbiome composition in *Formica*. Each bar depicts the relative abundance of ASVs in the microbiome of a single queen. ASVs have been grouped and coloured by genus. ASV’s that were found in only one sample or occurred at <10% relative abundance were collapsed into the *Other category. The full ASV table containing all bacterial genera is available in Supplementary Materials. The *Formica* host phylogeny is from Jackson et al. (Jackson et al., 2022), which combined a previous *Formica* phylogeny (Romiguier et al., 2018) with new sequencing information from Jackson et al. (2022).

As previously reported by Jackson et al. (2022), we found that *Liliensternia* is restricted to a single clade of *Serviformica* ants, while *Wolbachia* is present in every species screened and the majority of queens screened overall (112/131). *Wolbachia* is not only widely distributed among *Formica* species but also highly abundant, accounting for 54% of all sequences across all queens analysed. However, while most of the sequences came from *Wolbachia* in many queens, this was not the case in queens that were also infected with *Liliensternia*. When queens were co-infected with *Liliensternia* and *Wolbachia,* only a minority of the sequences typically came from *Wolbachia*.

Aside from *Wolbachia*, we found that two other bacterial genera commonly occur in *Formica* ants: *Lactobacillus* and *Spiroplasma. Lactobacillus* was taxonomically widespread, appearing in every clade of *Formica*. *Lactobacillus* occurred usually at low (< 25%) relative abundance, but in several *Serviformica* queens, it was the dominant bacterium. In contrast, *Spiroplasma* was relatively uncommon, being found in only five species (*F. rufibarbis*, *F. cinerea*, *F. selysi*, *F. lemani*, and *F. sanguinea*), where it typically occurred at < 25% relative abundance.

Some species harboured distinct microbial associates or microbiomes (Fig. 1). *Formica cinerea* was the only species recorded to carry *Serratia. Formica rufibarbis* was nearly always infected with *Spiroplasma*, although this latter host species was only collected at two sites. In *F. sanguinea*, there were two distinct microbiomes, as individuals had either *Wolbachia*-dominated microbiomes or microbiomes with *Spiroplasma*. The two distinct microbiomes of *F. sanguinea* correlated with two colonies in which this species was sampled in nature.

Most of the commonly occurring microbes in *Formica*, e.g. *Wolbachia*, *Liliensternia*, and *Spiroplasma*, typically have only a single copy of the 16s rRNA gene (Stoddard et al., 2015), so the abundance of these genera is unlikely to be a by-product of high gene copy number. In contrast, *Lactobacillus* can vary in copy number, ranging from 3 to 12 (Stoddard et al., 2015), so its abundance may be overestimated.

### Microbiome Composition across *Formica* Species

To test whether the microbiomes of *Formica* had distinguishable structure across species and clades, we analysed the composition of individual ASVs with permanova and beta-dispersion analyses. *Serviformica* clades 1 and 2 were analysed separately, as they are paraphyletic (Romiguier et al., 2018). We complemented the statistical analysis with a metaNMDS visualization, to aid in interpretation of the data (Fig. 2).

**Figure 2:**
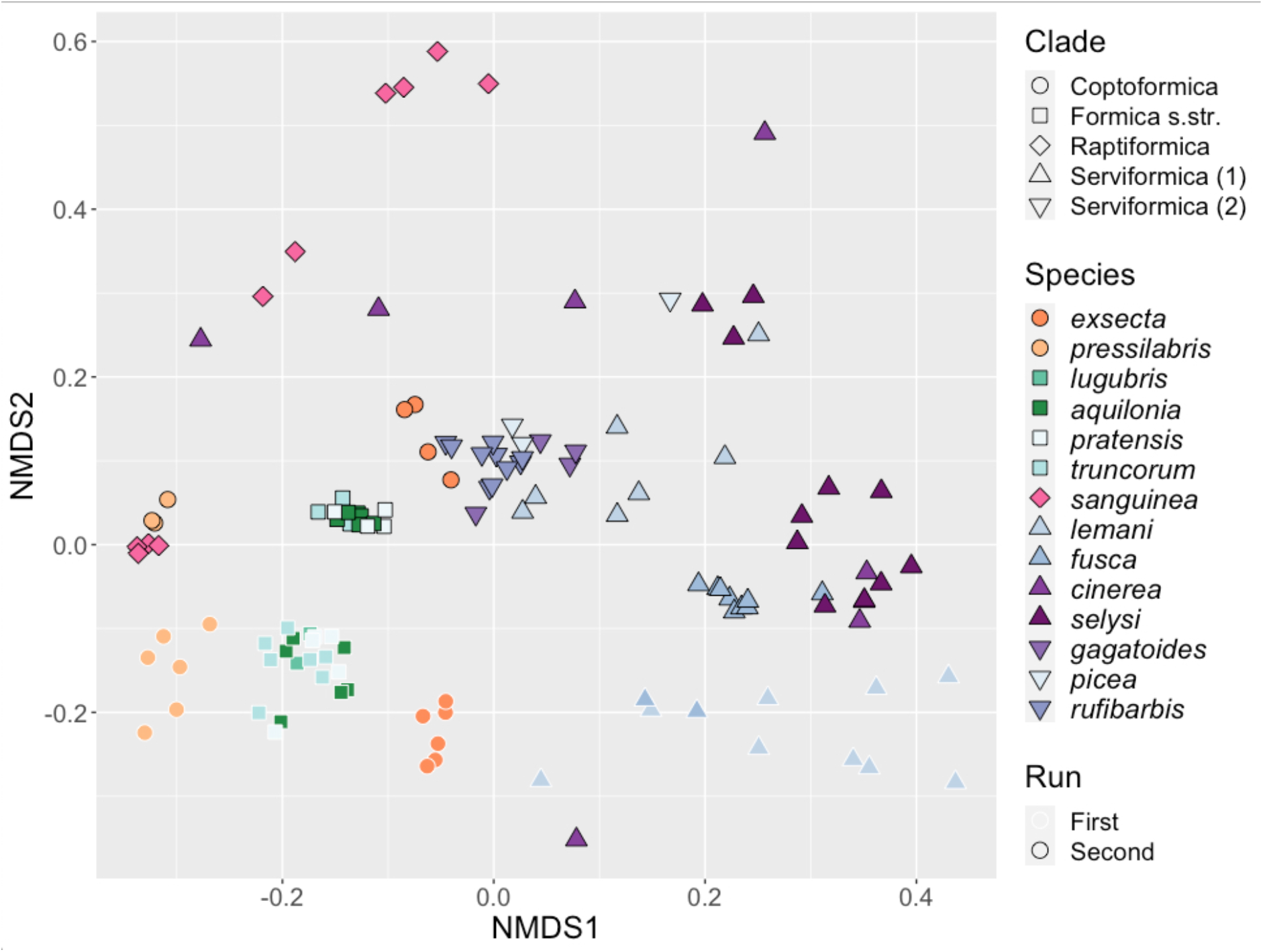
Similarity of microbial communities hosted by *Formica* queens. NMDS clustering of bacterial ASVs associated with individual *Formica* queens. Points clustering together represent samples with more similar microbial communities. Point shape designates clades, and colours represent different species. Points with a black outline are from the first run of 16S rRNA sequencing, points with a white outline are from the second run.

Our analysis revealed an effect of the sequencing run (Permanova: F1,128=17.889, P<0.001). However, we still found highly significant differences across host clades and species when we analyzed the runs separately (by blocking), or together with run as an interactive effect (All P<0.001, Table S2). We also found that there were significant differences in dispersion between taxonomic groups (Clade: F4,125=6.15, P<0.001, Species: F13,116=4.89, P<0.001). Individuals from some species and clade, such as *F. exsecta* and *F. pressilabris,* showed high degrees of similarity in their microbiome profile, resulting in low group-level dispersion. In contrast, other taxonomic groups, like *F. cinerea,* showed much higher levels of dissimilarity in microbiome structure and dispersion (Fig. 2).

We used a SIMPER test to investigate which ASVs were driving these differences between species and clades (Table S3). Ten bacterial genera accounted for more than 1% of the variation between *Formica* species: *Acinetobacter*, *Rhizobium*, *Lactobacillus*, *Pseudomonas*, *Serratia*, *Liliensternia*, *Spiroplasma*, *Stenotrophomonas*, *Tsukamurella*, and *Wolbachia*. Differences between *Formica* clades were explained by only four of these genera: *Lactobacillus*, *Liliensternia*, *Spiroplasma*, and *Wolbachia*.

In general, *Wolbachia* and *Liliensternia,* along with the two previously identified bacterial genera of interest, *Spiroplasma* and *Lactobacillus*, accounted for most of the variation between *Formica* species and clades. *Wolbachia* accounted for an average of 47±2% and 55±2% of the variation in microbial genera across host species and clades, respectively (Table S3). The other

ASV genus that accounted for a large amount of variation across species, but not clades, was *Liliensternia* (range of 10-67% across species, mean 33%). Despite being present in the majority of species and clades, *Lactobacillus* accounted for a small amount of the variation (range of 1-13% across species, mean 6%). Finally, *Spiroplasma* accounted for >1% of the bacterial variation across species within the clades *Serviformica* and *Raptiformica*. Other bacterial genera accounted for differences between a small number of ant species.

### Bacterial Ribotypes are Non-randomly Distributed across *Formica* Species

After identifying *Liliensternia*, *Wolbachia*, *Lactobacillus* and *Spiroplasma* as being the dominant bacterial genera composing *Formica* microbiomes (Fig. 2), we plotted the distribution of bacterial ribotypes across host species. The clustering of bacterial ASVs into 97% OTUs resulted in one *Liliensternia*, three *Wolbachia*, two *Spiroplasma*, and four *Lactobacillus* lineages (Fig. 3).

**Figure 3:**
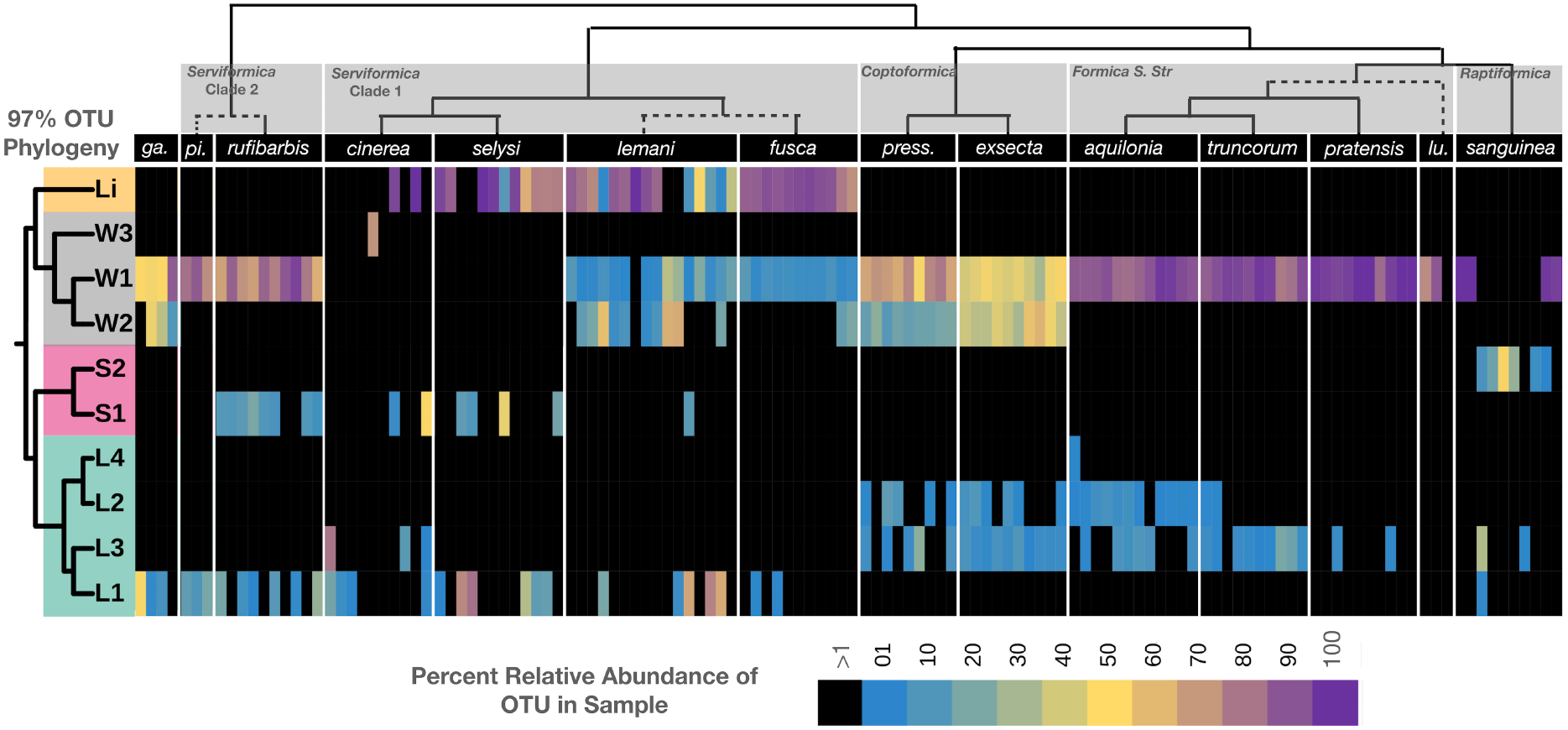
Bacterial ribotypes mapped on the phylogeny of *Formica* ants. The relative abundance of distinct lineage groups (based on 97% OTUs of the 16S rRNA gene) mapped on to the *Formica* phylogeny. Heatmap reflects the relative abundance of the four dominant bacterial genera found in each individual. Host and bacterial phylogeny are displayed above and to the left of the heatmap respectively. Prefix refer to: Li - *Liliensternia, W – Wolbachia*, S – *Spiroplasma*, and L - *Lactobacillus*.

The most dominant and widespread lineage was *Wolbachia* ribotype W1, which was found in every *Formica* species except *F. cinerea* and *F. selysi*, often at over 50% relative abundance. In contrast, *Wolbachia* ribotype W2 was found only in *Coptoformica* and *Serviformica* species. *Wolbachia* ribotype W2 was dominant in *F. exsecta,* whereas ribotype W3 appeared only in one *F. cinerea* individual.

The most widespread bacterial genus after *Wolbachia* was *Lactobacillus*. *Lactobacillus* Ribotype L2 was found exclusively in *Coptoformica* and *Formica S. Str.* species, while L3 was predominantly found in the same species as L2, plus *F. sanguinea*. *Lactobacillus* L1 was identified in species belonging to the *Serviformica* clades 1 and 2, as well as in one *F. sanguinea* individual, whereas L4 was only found in a single *F. aquilonia* individual.

There were two unique *Spiroplasma* ribotypes, one found in some *Serviformica* species and the other in *F. sanguinea*. A single ribotype associated with *Liliensternia* was found in one clade of *Serviformica*, as previously reported (Jackson et al., 2022), and in a single *F. pratensis* individual.

Within each bacterial genus, each ribotype tended to be associated with a restricted number of *Formica* clades, and in several cases more than one ribotype was present in the same individual. *Wolbachia* lineage W1 stood out from this pattern in that it appeared across all host clades, although it still often co-occurred with other ribotypes.

### Interactions between Symbiotic Bacterial Genera

We hypothesized that interactions between genera of bacteria may be one of the factors contributing to the overall structure observed in *Formica* microbiomes. Using ASV data from each amplicon sequencing run, we evaluated whether the presence of any genus of ASV was significantly associated with the presence or absence of another genus of ASV (Fig. 4, full statistics for all comparisons are available in Table S4). This analysis was intended to reveal co-occurrences between different bacterial genera. Further investigations will be required to determine the cause of positive or negative relationships.

**Figure 4:**
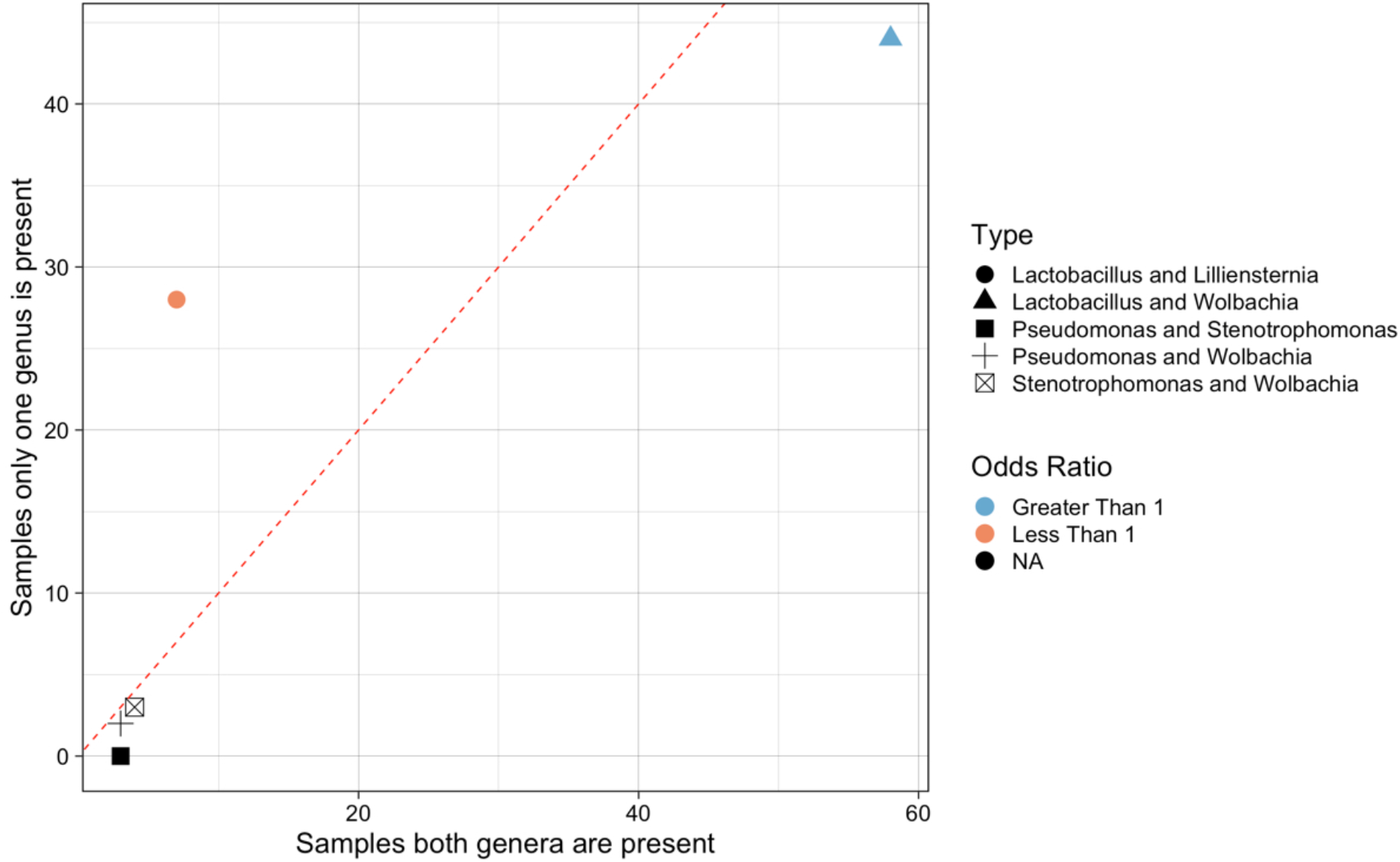
Significant positive and negative co-occurrences between microbes in *Formica*. Significantly co-occurring bacterial genera (under a Fisher’s exact test and FDR correction), by number of samples where both genera are present versus number of samples where only one of the two genera are present. Each microbe is represented by ASVs collapsed into taxonomic genus. Points are coloured by the odds ratio calculated by the Fisher test. Odds ratio’s greater than 1 (blue) indicate significant co-occurrence, where as ratios less than 1 (red) indicate a significant lack of co-occurrence. Odds ratios are not meaningful when one of the values is zero, these points are coloured black. Point shape denote different genera that shown significant correlation. Dotted line divide genera that tend to co-occur (below the line) from those that do not tend to co-occur (above the line). All data available in Supplementary Table S4.

We found five instances of significant co-occurrences between bacterial genera. *Lactobacillus* and *Liliensternia* were negatively correlated, i.e., they were less likely to co-occur than expected by chance. In contrast, *Lactobacillus* and *Wolbachia* tended to be positively associated, and commonly co-occurred (Fig. 3). There were also significant co-occurrences of *Pseudomonas* and *Stenophomonas*, *Pseudomonas* and *Wolbachia*, and *Wolbachia* and *Stenophomonas.* However, these last three co-occurrences should be viewed with caution as *Pseudomonas* and *Stenophomonas* only occurred in a small number of samples (<5 each), so there is limited power to resolve their associations with other microbes.

### Phylosymbiosis in *Formica* ants

Given the strong structure of microbial ASVs and ribotypes associated with different *Formica* species and clades, we tested whether microbiome composition mirrors the phylogeny of *Formica* species, i.e. phylosymbiosis. We also tested whether individual bacterial genera showed signs of phylogenetic structure across hosts. Using the protocol established by Brooks et al. (2017), we generated trees based on the differences in abundance of each unique ASV across the microbiome of each *Formica* species (Fig. 5), and compared these to the host phylogeny using Robinson-Folds clustering, as well as unadjusted and normalized Matching Cluster metrics.

**Figure 5:**
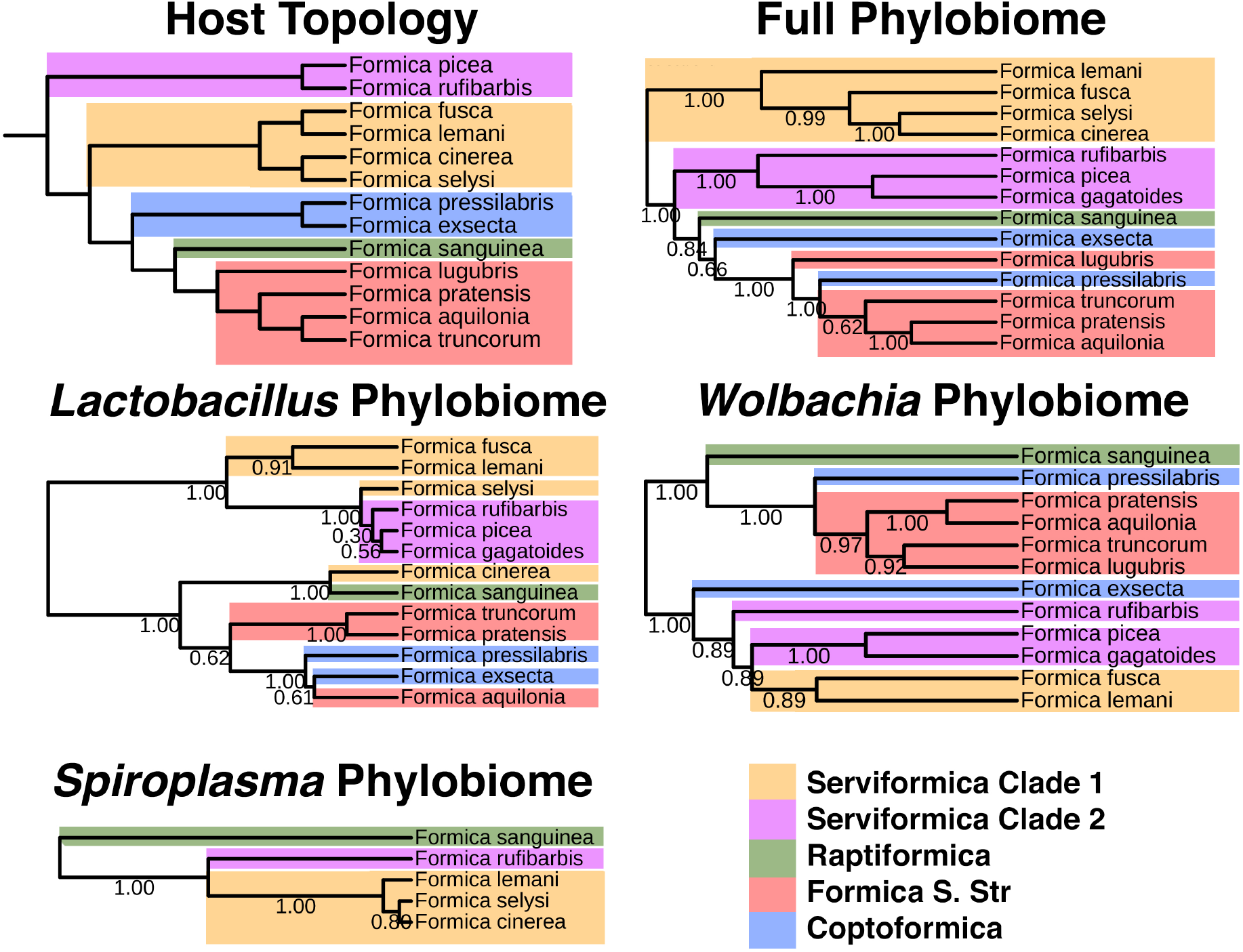
Host phylogeny versus Phylobiomic signatures of *Formica* species. Bootstrap values on bacterial phylogenies indicate proportion out of 100 iterations of subsampling within microbiome data resulted in the given placement of branches. Bacterial phylogeny generated using the protocol established by Brooks et al. (2017).

All phylobiome trees were highly congruent with the host tree (P<0.01; Table S5). In addition to significantly corresponding to the host tree, we documented two interesting patterns when comparing phylobiome trees (Fig. 5). First, the paraphyletic nature of *Serviformica* species was mirrored in all phylobiome trees. We expected this pattern in the full phylobiome tree, because one clade of *Serviformica* carries *Liliensternia*, a strictly vertically transmitted symbiont whose phylogeny mirrors that of its host (Jackson et al., 2022). However, even when we considered trees of the single bacterial genera *Spiroplasma, Wolbachia,* and *Lactobacillus,* the paraphyletic pattern was still apparent (Fig. 5). Second, the position of *Coptoformica* species (*F. exsecta* and *F. pressilabris*) was not consistent across phylobiome trees (Fig. 5).

## Discussion

### Components of a Highly Structured Microbiome in Formica Ants

Our investigation has revealed that *Formica* ants contain highly structured stable microbiomes that recapitulate the phylogeny of their host. The structure is principally due to the prevalence and distribution of four genera of bacteria: *Wolbachia*, *Lactobacillus*, *Liliensternia*, and *Spiroplasma*. Of the four, *Wolbachia*, *Liliensternia,* and *Lactobacillus* have been previously identified as symbionts of *Formica* (Russell, 2012; Jackson et al., 2022; Zheng et al., 2022), whereas we have newly identified *Spiroplasma* as a common microbe associated with several *Formica* species, suggesting it may be symbiotic in nature. The most dominant microbes in terms of relative abundance were *Wolbachia* and *Liliensternia*, a pattern likely explained by the fact that we analysed adult queens. Queens have large reproductive tissues, in which *Wolbachia* and *Liliensternia* are maintained (Moran et al., 2008; Frost et al., 2014; Jackson et al., 2022).

*Wolbachia*, the most abundant bacterium across *Formica* queens, is a common heritable reproductive manipulator found in a wide range of insects (Zug and Hammerstein, 2012). It is also found in many ant species, although it is unclear to what extent it behaves as a reproductive manipulator in ants (Keller et al., 2001; Russell, 2012). The frequency of *Wolbachia* infection within *Formica* species is particularly high compared to other ants and insects (Russell, 2012). Our results are consistent with previous measurements of *Wolbachia* infection in *Formica* species, which indicated that over 80% of individuals were infected (Russell, 2012; Russell et al., 2012). In *F. truncorum*, *Wolbachia* is close to fixation, and heavily infected colonies produced significantly fewer sexuals, suggesting the microbe has a deleterious effect in this species, despite its prevalence (Wenseleers et al., 2002). One factor that has been proposed to contribute to the varying success of *Wolbachia* in ants is variation in effective population size, where species with small effective population sizes resulting from limited queen dispersal or dependent colony founding have higher rates of *Wolbachia* infection (Wenseleers et al., 1998; Treanor and Hughes, 2019). However, we observe high rates of *Wolbachia* infection across all *Formica* species, which included those with different colony founding strategies, queen dispersal strategies, and lifestyles.

*Lactobacillus* has been described in multiple ant species (Anderson et al., 2012), including *F. exsecta* (Johansson et al., 2013; Zheng et al., 2022). The contribution of *Lactobacillus* to the structured microbiomes of *Formica* ants pairs well with recent findings that ants that frequently feed on aphid honeydew, such as *Formica*, commonly carry lactic acid bacteria that help catabolize sugars known to be found in the sugary excretion (Engel et al., 2012; Zheng et al., 2022). However, it was surprising that it appeared to contribute to inter-clade and inter-species differentiation of *Formica* microbiomes. *Lactobacillus* is typically a gut-associated microbe (Zheng et al., 2022) and is not known to be ovarially transmitted. As a result, phylogenetic signal deriving from *Lactobacillus* strains is not likely to be driven by inheritance, unless long term vertical transmission has been sustained through trophallaxis. Additionally, it is not expected for all ants to possess stable gut microbiomes, as many species appear to exist devoid of an appreciable gut bacterial community (Sanders et al., 2017). Studies have shown that *Lactobacillus* can serve as a defensive or nutritional symbiont in *Drosophila* and in honeybees (Forsgren et al., 2010; Storelli et al., 2011; Vásquez et al., 2012). If *Lactobacillus* served a similar role in *Formica,* it might explain why it has established a stable presence across species, as previously proposed by Zheng et al. (2022).

*Spiroplasma* is a common heritable insect endosymbiont that is often pathogenic and known for its ability to manipulate the reproduction of its host (Majerus et al., 1999; Jiggins et al., 2000). However, studies have also shown that *Spiroplasma* can benefit insects, by defending against pathogens such as fungi (Łukasik et al., 2013). *Spiroplasma* has been detected in several ant species (Ishak et al., 2011; Ballinger et al., 2018). In *Solenopsis* and *Myrmica* ants, *Spiroplasma* is found in nearly all individuals, suggesting it may play a beneficial role for its host (Ishak et al., 2011; Ballinger et al., 2018). The role of *Spiroplasma* in *Formica* ants is currently unclear. As *Spiroplasma* was very common in *F. rufibarbis* and *F. sanguinea* - found at high frequencies in 10 colonies from 5 sites across Finland - it would be interesting to know whether a parasitic, or potentially a beneficial relationship, explains its prevalence in these two host species.

Previous studies have shown that *Liliensternia* is housed in bacteriocytes that surround the midgut (Lilienstern, 1932; Jackson et al., 2022), and is strictly vertically inherited from the common ancestor in *Serviformica* clade 1(Lilienstern, 1932; Jackson et al., 2020; Jackson et al., 2022). This explains the overwhelming presence of *Liliensternia* in this clade of ants. Normally, a bacteriocyte-associated symbiont would be present in all reproductive females. However, the curious feature that *Liliensternia* is being lost in some queens, especially in species such as *F. cinerea* (Jackson et al., 2022), explains why it is not universally found in queens within this clade of hosts.

### Synergy and Antagonism in the Microbiome

Our analysis of pairs of bacterial genera across *Formica* microbiomes revealed both positive and negative associations between microbes. Some of these correlations indicated that there may be some underlying synergism or antagonism between microbes. For example, we found a positive relationship between *Wolbachia* and *Lactobacillus*. This was unexpected, as these two symbionts are likely localized in different parts of the insect and have different routes of transmission - ovarial versus environmental -, so there would be fewer opportunities for the microbes to interact. It may be that an indirect interaction mediated through the host leads to the positive association. However, at present it is unclear how this type of interaction would occur. We also found that *Liliensternia* does not tend to co-occur with *Lactobacillus*. This was intriguing, as *Lactobacillus* is known to behave as a nutritional mutualist in some other insect species (Storelli et al., 2011). It may be that *Lactobacillus* does not tend to co-occur with *Liliensternia* because the host only needs one symbiont to take on the nutrient provisioning role.

### Phylosymbiosis in *Formica* Ants

Our analyses revealed that *Formica* microbiomes show evidence of phylosymbiosis, in that the bacterial community composition mirrors the phylogeny of the host (Lim and Bordenstein, 2020). A wide range of mechanisms can lead to phylosymbiosis, including vertical transmission of microbes, interactions between microbes, host-microbe genetic compatibility, or the similarities in host ecologies that impact the microbes they carry (e.g. diets). In *Formica* ants, several life history traits correlate with host phylogeny (Borowiec et al., 2020), including colony founding method and mound building style. As a result, it is not possible to fully disentangle to which extent similarities in the ecologies, or the phylogenetic proximity of *Formica* species, contribute to the microbes they carry. However, our results suggest that host phylogenetic relatedness, in combination with vertical transmission, may play important roles.

We have shown that *Formica* microbiomes are largely dominated by vertically transmitted microbes. This includes *Liliensternia,* which is believed to be an ancient strictly vertically transmitted symbiont that has co-speciated with its host (Jackson et al., 2022), and undoubtedly contributes to the phylosymbiosis signal. However, even when *Liliensternia* was not considered in the analysis - as is the case in the *Lactobacillus*, *Wolbachia*, and *Spiroplasma* phylobiome trees - *Formica* still show signals of phylosymbiosis, demonstrating that the pattern isn’t solely driven by the presence of this nutritional mutualist. Phylogenetic studies of *Wolbachia* and *Spiroplasma* have shown that although both microbes are maternally transmitted, they are often horizontally transferred between host lineages on evolutionary time scales, and therefore typically do not co-speciate with their hosts (e.g. (Viljakainen et al., 2008; Ahmed et al., 2016). There are a few instances where co-diversification has been observed, for example between *Spiroplama* and *Myrmica* ants, and between *Wolbachia* in both filarial nematodes and bedbugs (Fenn and Blaxter, 2004; Ballinger et al., 2018; Balvín et al., 2018). The widespread occurrence of the same *Wolbachia* and *Spiroplama* ribotypes (e.g. W1 & S1) in distantly related *Formica* species found in our data, and in a separate study on *Wolbachia* in *Formica* (Viljakainen et al., 2008), indicates that horizontal transfer between species is relatively common on evolutionary time scales. Therefore, vertical inheritance from a common ancestor is unlikely to have led to the observed pattern of phylosymbiosis. The high frequency of specific *Wolbachia* strains in certain *Formica* species is therefore more likely the product of host specific factors, e.g. genetic compatibilities of *Wolbachia* strains with related hosts increasing the likelihood that the symbiont is horizontally transferred between related species, with vertical transmission helping to maintain the relationship within populations and species.

The life-history traits of *Formica* also provide some capacity to tease apart the importance of host ecology and phylogeny in structuring their microbiomes. All of the *Formica* ant species studied, except those in the *Serviformica* clades, have lost the ability to found their colonies independently, and rely partly or completely on socially parasitic strategies for colony founding (Romiguier et al., 2018; Borowiec et al., 2021). This means they take over colonies of other ants, specifically those in the *Serviformica* clades, to found new colonies (temporary social parasitism). Additionally, *F. sanguinea* facultatively raids workers from *Serviformica* species to complement their own worker force. As a result, different clades of *Formica* ants come into close association with each other regularly. However, despite coming into close physical proximity with other species - living within the same colony and feeding other species via trophallaxis - we still find that socially parasitic species and the species they parasitize have distinct microbial profiles. This indicates that environmental acquisition or transfer between unrelated species in close physical proximity does not play a significant role in structuring the microbiomes of *Formica* ants. Given this, we hypothesize that phylogenetic relatedness of *Formica* species plays a more substantive role in the retention of microbial species, and the overall similarity of microbiomes between related host species found in our study. Studies have shown that the horizontal transfer of heritable intracellular symbionts is more likely to occur between closely related species (Łukasik et al., 2015; McLean et al., 2019). Furthermore, recent studies on aphids have shown that co-adaptation of facultative symbionts to different hosts, and their specific ecologies, is largely responsible for the distribution of symbiont genotypes across host species (Henry et al., 2022; Wu et al., 2022) However, we cannot exclude that similarities in host ecology may also play a role in the phylosymbiotic signal we find in *Formica.* Recent studies on *Formica* and *Lasius* ants demonstrated that *Lactobacillus* was more common in ants that feed on aphid honeydew (Zheng et al., 2022). *Formica* ants commonly tend aphids for honeydew and species may differ in their reliance on aphids for this resource (Fiedler et al., 2007). It is possible that the prevalence of *Lactobacillus*, and its contribution to the phylosymbiotic signal may be partially explained by shared dietary ecologies, if for example related *Formica* species have similar tendencies to feed on honeydew as a food source.

### Conclusions

We have shown that *Formica* ants have formed stable enduring relationships with a core set of microbes that are dominated by four genera of bacteria. When placed in a phylogenetic context, we reveal that the composition of microbes carried by different *Formica* species recapitulates the phylogeny of the host. For the bacteriocyte-associated *Liliensternia*, it is believed that the association between host and microbe is perpetuated by a mutualistic benefit, with vertical transmission explaining its occurrence in related species. For the other dominant microbes, we hypothesize that the genetic proximity among microbes and among hosts, respectively, and possibly host feeding ecology, may explain the communities of microbes associated with *Formica* ants. The relationship with *Lactobacillus* and *Formica* may also be mutualistic in nature, by helping their host digest honeydew (Engel et al., 2012; Zheng et al., 2022), which is an important food source for many *Formica* species. It is less clear, however, why many *Formica* species share a strong, near ubiquitous, association with *Wolbachia*. Future studies should investigate whether *Wolbachia*’s high prevalence in *Formica* reflects a very successful pathogen, or whether the ants have evolved some form of dependence on the microbe (Comandatore et al., 2015), which may help explain its prevalence across host species.

## Data Availability

All data collected in association with this paper are available under bioproject accession PRJNA639935.

## Competing Interest Statement

We declare no conflict of interest.

## Acknowledgments

The authors thank Dominic Burns for providing ant samples. This project was funded by L.M.H.’s NERC IRF (NE/M018016/1).

## Notes

https://www.ncbi.nlm.nih.gov/bioproject/?term=PRJNA639935

### Competing Interest Statement

The authors have declared no competing interest.

